# Relational integration demands are tracked by temporally delayed neural representations in alpha and beta rhythms within higher-order cortical networks

**DOI:** 10.1101/2024.10.16.618779

**Authors:** Conor Robinson, Luca Cocchi, Takuya Ito, Luke Hearne

## Abstract

Relational reasoning is the ability to infer and understand the relations between multiple elements. In humans, this ability supports higher cognitive functions and is linked to fluid intelligence. Relational complexity (RC) is a cognitive framework that offers a generalisable method for classifying the complexity of reasoning problems. To date, increased RC has been linked to static patterns of brain activity supported by the frontoparietal system, but limited work has assessed the multivariate spatiotemporal dynamics that code for RC. To address this, we conducted representational similarity analysis in two independent neuroimaging datasets (Dataset 1 fMRI, n=40; Dataset 2 EEG, n=45), where brain activity was recorded while participants completed a visuospatial reasoning task that included different levels of RC (Latin Square Task). Our findings revealed that, spatially, RC representations were widespread, peaking in brain networks associated with higher-order cognition (frontoparietal, dorsal-attention, and cingulo-opercular). Temporally, RC was represented in the 2.5 - 4.1 seconds post-stimuli window and emerged in the alpha and beta frequency range. Finally, multimodal fusion analysis demonstrated that shared variability within EEG-fMRI signals within higher-order cortical networks were better explained by the theorised RC model, relative to a model of cognitive effort (CE). Altogether, the results further our understanding of the neural representations supporting relational processing, highlight the spatially distributed coding of RC and CE across cortical networks, and emphasise the importance of late-stage, frequency-specific neural dynamics in resolving RC.

## Introduction

Relational reasoning, the ability to mentally manipulate and integrate relationships between interacting elements, is a core component of higher-order cognition and a hallmark of human intelligence (Waltz et al., 1998; Halford et al., 2010). Studies on relational reasoning reveal fundamental capacity limitations in the number of relations that can be processed simultaneously (Halford et al., 2005). This constraint is formalised within relational complexity (RC) theory, a framework that has uncovered performance boundaries in everyday problem-solving (Boag et al., 2006), neurodevelopment (Halford et al., 1998; Eslinger et al., 2009; Todd et al., 2019), and clinical populations (Goel and Grafman, 1995; Hearne et al., 2019). While numerous studies have demonstrated the broad utility of RC theory, a key open question remains: where and when does the brain represent RC?

Functional magnetic resonance imaging (fMRI) work has provided a map of the brain *spatial* correlates of relational reasoning (Wertheim and Ragni, 2018). This body of work has led to the perspective that a frontoparietal network (FPN), also known as the multiple-demand network (MDN), is essential for performing relational reasoning tasks (Duncan and Owen, 2000; Cocchi et al., 2013; Hearne et al., 2017; Holyoak and Monti, 2021; Morin et al., 2022). Further work has emphasised interactions between the FPN and other brain systems, such as the cingulo-opercular and sensory networks while performing complex reasoning tasks (Cocchi et al., 2013; Cole et al., 2013; Hearne et al., 2017). Studies have typically emphasised the univariate association between specific brain activations (or functional connections) and relational demands, rather than how a given brain region might code for reasoning complexity (Freund et al., 2021). Thus, it remains unclear how the spatial correlates described by prior work relate to neural representations underlying RC. It is also plausible that the observed activations in the MDN might code for general cognitive effort (CE) rather than RC per se, as the MDN has been shown to track task difficulty even when RC is held constant (Wen and Egner, 2023). This underscores the need to determine whether patterns of neural activity specifically track RC or reflect general CE as relational demands increase.

Despite significant advances in the spatial mapping, the temporal dynamics of RC have not been well documented. A handful of electroencephalography (EEG) event-related studies have investigated reasoning demands, typically reporting deviations in the amplitude of event-related potentials as a function of complexity (Cutmore et al., 2015; Xiao et al., 2022). However, these studies did not analyse the entire cognitive processing period or frequency-resolved activity, largely due to uncertainties about how relational processing unfolds in time. While distinct from RC (Halford et al., 1998; Andrews et al., 2006), research on working memory in humans and non-human primates has demonstrated the crucial role of synchronised neural oscillatory activity in alpha and beta frequency bands (Miller et al., 2018; John et al., 2022; Fulvio et al., 2024), with high-frequency gamma (> 40 Hz) suggested to update sensory content (Honkanen et al., 2015; Lundqvist et al., 2018). Although working memory and relational complexity pose distinct cognitive challenges (Halford et al., 1998; Andrews et al., 2006), there may be overlap in their underlying neurophysiological mechanisms, particularly in how neural activity in alpha and beta frequency bands support the integration demands required for relational reasoning. Combining fMRI mapping with an analysis of the oscillatory temporal dynamics underpinning RC or CE is likely to assist in the delineation of a core temporal and spatial signature supporting the neural representation of abstract relational integration.

To explore the spatiotemporal representation of RC, we used the Latin Square Task (LST), a visuospatial reasoning task where participants solve short Sudoku-like puzzles (Birney et al., 2006; Hearne et al., 2020). Cognitive demands were modulated by changing the minimum number of relations that needed to be integrated to infer the solution. Representational Similarity Analysis (RSA) provided a means to fuse multiple datasets rich in multimodal spatial and temporal information to assess where and when RC is represented in the brain (Kriegeskorte et al., 2008). To this end, we pooled data from two independent studies assessing brain activity using fMRI and EEG while participants completed the LST at three levels of relational complexity. In line with prior studies, we predicted that RC representations would emerge in higher-order associative networks responsible for executive control in relational integration (Cocchi et al., 2013). We anticipated that the emergence of a RC representation across distinct brain systems would be observable in alpha, beta and gamma neural oscillation patterns. To distinguish RC from general task demands, we compared these representations against an alternative CE model. Finally, we extended our analysis by employing a model-based fusion approach (Cichy et al., 2014) to quantify the unique contribution of RC and CE to the shared spatiotemporal representation between neuroimaging datasets.

## Materials and Methods

### Participants

Participants were pooled across two independent datasets. In the first dataset, sixty-five healthy, right-handed participants completed the LST while undergoing fMRI (final n = 40, see exclusion criteria in the *fMRI preprocessing* section below; mean = 23 years old; age range = 18 - 33; 60% female). This dataset has been used in prior publications (Hearne et al., 2017; Shine et al., 2019). In the second dataset, forty-seven healthy participants completed the LST concurrently with EEG (final n=45, see the *EEG preprocessing* section for exclusions; mean = 25; age range = 19 - 33; 62% female). Ethics approval was granted by the University of Queensland Human Research Ethics Committee (dataset one) and the QIMR Berghofer Research Ethics Committee (dataset two). In both experiments, participants provided written informed consent and were eligible if aged between 18 - 35 with no previous reported history of a mental health or neurological disorder.

### Experimental design

The LST is a nonverbal relational reasoning experimental task (Birney et al., 2006). Each LST puzzle involves the presentation of a four-by-four grid populated by shapes (squares, triangles, circles and crosses) and a target question mark. Participants are asked to solve what shape the question mark must be, given that *an item or object can only exist once in every row and column* (similar to sudoku). Puzzle complexity (Binary, Ternary and Quaternary) was modulated by modifying the number of items that needed to be interrelated in one step to solve the problem. As illustrated in **Figure 1a**:

- Binary relations: required linking items across a single row or column,
- Ternary relations: required consideration across one row and one column, and
- Quaternary relations: required integration across multiple rows and columns.

**Figure 1:**
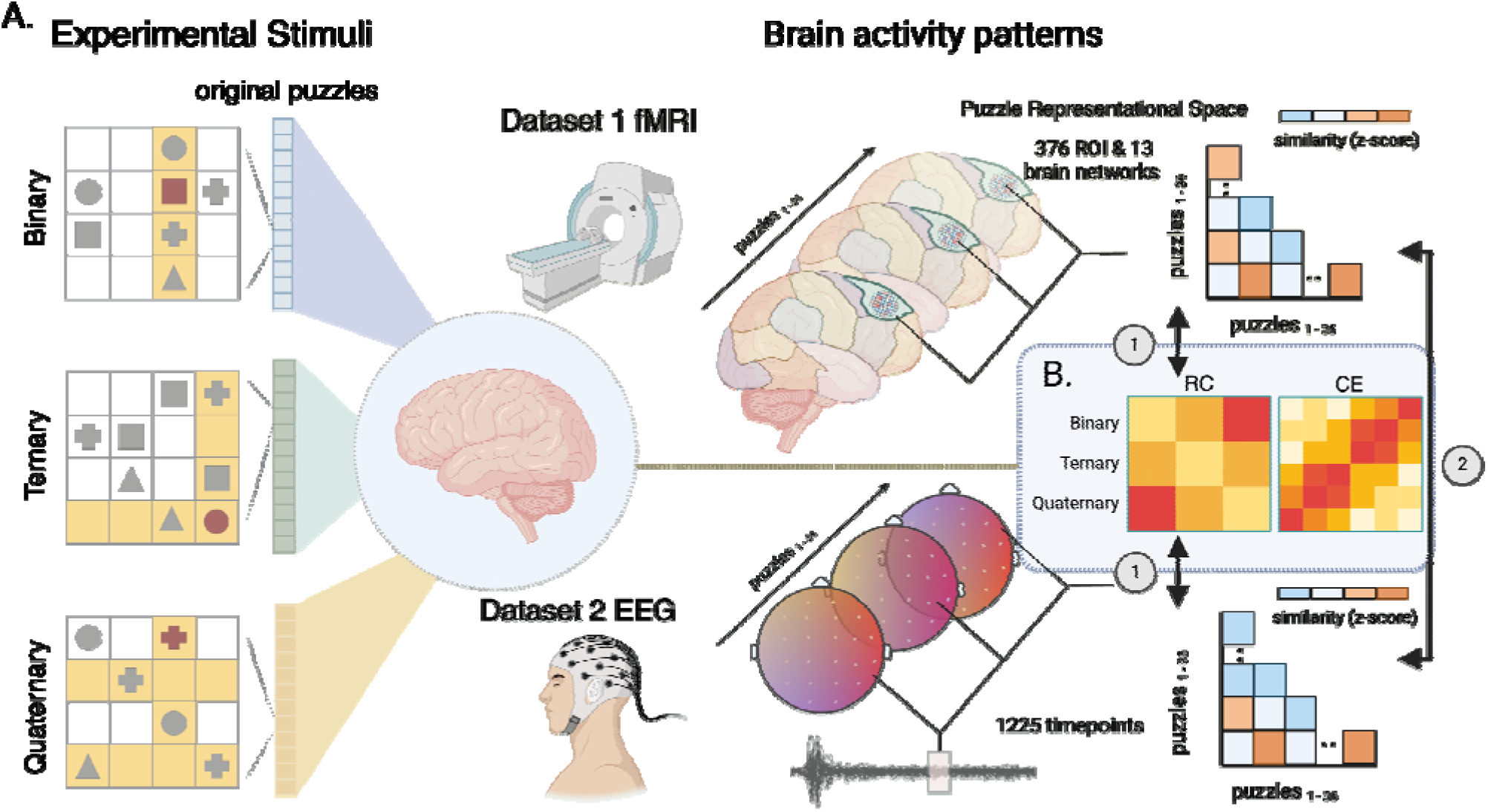
*Stimulus design and multivariate analysis approach*. Latin Square problems shown on the left required participants solve for the target square (solution highlighted in red) adhering to the rule that each item could only appear once in each row and column. Relational complexity was manipulated by varying the minimum number of interrelating rows and columns (highlighted in yellow) to solve for the missing item. The three puzzles illustrate example Binary, Ternary, and Quaternary complexity levels. Each cell in the corresponding-coloured vector represents one of the 36 original puzzle designs, with 12 puzzles per complexity level. While completing the task, brain activity was recorded using fMRI (dataset 1, top, gold) and EEG (dataset 2, bottom, light green). Neurophysiological responses were converted into puzzle representational dissimilarity matrices (RDMs) based on the distances between activation patterns evoked by different puzzles. Spatial fMRI RDMs were derived by correlating vertex or voxel activity patterns within parcellated brain regions for each puzzle. Temporal EEG RDMs were generated by correlating sensor-level activity patterns at each timepoint across puzzles. These spatial and temporal RDMs were (1) compared to the theoretical relational complexity (RC) model and data-driven cognitive effort (CE) model (B). Moreover (2) assessed for spatial- temporal co-occurrence.

Importantly, all LST problems across both studies are designed to be solvable in a single step, ensuring that working memory capacity was not differentially taxed across complexity levels. Rather, the increased processing load stemmed from the requirement to relate information from different sources across the grid. In the second dataset, participants solved identical puzzles, with the exception that different coloured circles replaced the geometric shapes used in the first study as visual stimuli (A, B, C and D).

Each relational complexity condition comprised of twelve unique puzzle designs. To increase the number of trials per condition, each of the 12 unique puzzle designs were rotated 90 degrees multiple times (Hearne et al., 2020; Hartung et al., 2022). In addition to the rotational transformations, the items occupying each position were pseudo randomised across participants. This item randomisation ensured that in the unlikely situation that a participant recognised a rotated puzzle, they would need to perform relational reasoning to solve the problem, as the specific item-position mappings were unique for each presentation. Together, this created new puzzle configurations while preserving the relational topology of the 12 original problems. In the first dataset, this was performed twice to generate 36 trials per complexity level. In the second dataset, this was performed thrice to generate 48 trials at each complexity level. Increased trial numbers were warranted in the second dataset due to EEG artefact rejection. For each dataset, the puzzle rotations were averaged to generate brain activation estimates for twelve trials per condition, thirty-six trials in total (see the *representation similarity analysis* section below for details).

Trials were presented in a pseudo randomised order, ensuring that two trials of the same complexity level were not presented sequentially. To optimise the number of trials for imaging acquisition the stimulus duration was slightly modified for each modality. In the first dataset, problems were presented for 5 seconds, with a further 2 seconds to respond via a response screen. In the second dataset, the stimulus time was 4.5 seconds puzzle onset and a 1.75 second response window.

### Neuroimaging acquisition, preprocessing and activation estimation

#### fMRI Dataset

As reported previously (Hearne et al., 2017), fMRI data were acquired on a 7 Tesla Siemens scanner equipped with a 32-channel head coil at the Centre of Advanced Imaging, The University of Queensland, Brisbane. Whole-brain echo-planar images were collected with the following parameters: isotropic 2 mm voxels, TR = 586ms, TE = 23ms, flip angle = 40°, FOV = 208mm, 55 axial slices, multiband acceleration factor = 5. The LST was performed over three 12-minute runs (1250 volumes). Structural scans (MP2RAGE) used in the preprocessing pipeline were acquired with the following parameters: isotropic 0.75 mm voxels, TR = 4300ms, TE = 3.44 ms, and 256 slices. Stimuli were presented using PsychToolBox 3.0 (Brainard, 1997) onto a screen located at the head end of the MR scanner, and participants viewed the projected stimuli via a mirror head-mount. The grid layout of the puzzle was sized to fit within the field of view of participants, removing the need for large eye-moment to peruse the grid space.

Data were preprocessed using fMRIPrep 21.0.1 (Esteban et al., 2019) [RRID:SCR_016216], which is based on *Nipype* 1.6.1 (Gorgolewski et al., 2011) [RRID:SCR_002502]. Due to differences in signal intensity in the temporal cortex with the rest of the brain, the raw T1- weighted (T1w) images were intensity corrected using the SPM segmentation tool prior to the fMRIPrep pipeline. Based on quality control checks this step ensured that different tissues were correctly identified during the preprocessing pipeline. Following this, within the fMRIPrep pipeline, the T1w image was corrected for intensity non-uniformity (INU) with N4BiasFieldCorrection (Tustison et al., 2010), distributed with ANTs 2.3.3 (Avants et al., 2008) [RRID:SCR_004757], and used as T1w-reference throughout the workflow. The T1w-reference was then skull-stripped with a *Nipype* implementation of the antsBrainExtraction.sh workflow (from ANTs), using OASIS30ANTs as target template. Brain tissue segmentation of cerebrospinal fluid (CSF), white-matter (WM) and gray-matter (GM) was performed on the brain-extracted T1w using fast (FSL 6.0.5.1:57b01774, RRID:SCR_002823, (Zhang et al., 2001)). Brain surfaces were reconstructed using recon-all (FreeSurfer 6.0.1, RRID:SCR_001847, (Dale et al., 1999)), and the brain mask estimated previously was refined with a custom variation of the method to reconcile ANTs-derived and FreeSurfer-derived segmentations of the cortical gray-matter of Mindboggle (RRID:SCR_002438, (Klein et al., 2017)). Volume-based spatial normalization to standard space was performed through nonlinear registration with antsRegistration (ANTs 2.3.3), using brain-extracted versions of both T1w reference and the T1w template. *FSL’s MNI ICBM 152 non-linear 6th Generation Asymmetric Average Brain Stereotaxic Registration Model* was used for spatial normalisation [(Evans et al., 2012), RRID:SCR_002823; TemplateFlow ID: MNI152NLin6Asym].

For each of the three BOLD runs found per subject the following preprocessing was performed. First, a reference volume and its skull-stripped version were generated by aligning and averaging 1 single-band references (SBRefs). Head-motion parameters with respect to the BOLD reference (transformation matrices, and six corresponding rotation and translation parameters) are estimated before any spatiotemporal filtering using mcflirt (FSL 6.0.5.1:57b01774, (Jenkinson et al., 2002)). BOLD runs were slice-time corrected to 0.268s (0.5 of slice acquisition range 0s- 0.535s) using 3dTshift from AFNI ((Cox and Hyde, 1997), RRID:SCR_005927). The BOLD time-series were resampled onto their original, native space by applying the transforms to correct for head-motion. These resampled BOLD time-series will be referred to as *preprocessed BOLD in original space*, or just *preprocessed BOLD*. The BOLD reference was then co-registered to the T1w reference using bbregister (FreeSurfer) which implements boundary-based registration (Greve and Fischl, 2009). Co-registration was configured with six degrees of freedom. First, a reference volume and its skull-stripped version were generated using a custom methodology of *fMRIPrep*. The three signals are extracted within the CSF, the WM, and the whole-brain masks. The BOLD time-series were resampled onto *fsaverage* surface space. *Grayordinates* files (Glasser et al., 2013) containing 91k samples were also generated using the highest-resolution fsaverage as intermediate standardized surface space. All resamplings can be performed with *a single interpolation step* by composing all the pertinent transformations. Gridded (volumetric) resamplings were performed using antsApplyTransforms (ANTs), configured with Lanczos interpolation to minimize the smoothing effects of other kernels (Lanczos, 1964). Non-gridded (surface) resamplings were performed using mri_vol2surf (FreeSurfer). Given the RSA calculates similarities between voxel / vertex activation patterns without spatial smoothing, any subjects exceeding 2 mm of head motion (six rotation and translation parameters) were excluded from further analysis (final *n*=40). After fMRIPrep, the functional data were further denoised by regressing fourteen nuisance signals (mean cerebrospinal fluid and white matter signals, along with the six motion parameters and their derivatives) using Nilearn (Abraham et al., 2014).

To estimate brain activations, whole-brain general linear model (GLM) analyses were conducted using Nilearn (Abraham et al., 2014) for each participant’s grayordinate surface and volumetric data. Using a previously established method to estimate trial-by-trial activation patterns (Mumford et al., 2012), a separate GLM was performed for each of the 36 puzzles in the experiment. Each design matrix included two regressors; one for the three trials of interest (three rotations of the puzzle) and another for all other trials in the experiment. Each trial was modelled as a five second boxcar regressor convolved with the canonical hemodynamic response function. Three additional regressors were included to account for each run of the experiment. This procedure resulted in brain activation estimates for each of the 36 puzzles across the brain which were subsequently analysed using RSA.

#### EEG Dataset

EEG recordings were digitised at 2000 Hz using an ANT-neuro system with a 64 channel cap arranged in the 10-20 layout. Experimental stimuli were presented on a BenQ 24 inch computer monitor with the task paradigm implemented using PsycoPy toolbox (v2020.2.10) (Peirce et al., 2019). Similar to Dataset 1, the grid layout of the puzzle was sized to fit within the field of view of participants.

EEG recordings were preprocessed offline using EEGLAB (Delorme and Makeig, 2004). The continuous data underwent high-pass finite impulse response (FIR) filtering at 0.5 Hz to remove low-frequency drifts. Line noise centred at 50 Hz was eliminated using a notch filter spanning 47.5-52.5 Hz, followed by a 97 Hz low-pass filter to attenuate harmonic components. Bad channels, identified through visual inspection, were excluded before re-referencing to the average of all remaining channels, excluding the M1, M2, and EOG electrodes. The data were then downsampled to 250 Hz. Trial epochs were generated using the stimulus onset as a marker resulting in a -400 ms pre-stimulus and 4500 ms post-stimulus window. After independent component analysis (ICA) based artefact rejection, the excluded bad channel data were reconstructed using spherical spline interpolation. Finally, the signal underwent band-pass filtering within the EEG observable range of 1-45 Hz. Two subjects were removed from the final analysis due to excessive muscle artefacts.

EEG data was further processed and frequency decomposition performed using the MNE Python library (Gramfort et al., 2013). Epochs were baseline-corrected by subtracting the mean signal during the pre-stimulus period from the entire epoch (stimuli onset to 4500 ms). To investigate oscillatory dynamics across different frequency bands, the epoched timeseries data were decomposed into canonical frequency bands using zero-phase bandpass filtering. Specifically, the following frequency ranges were extracted: theta (4-8 Hz), alpha (8-12 Hz), beta (13-30 Hz), and gamma (30-45 Hz).

### Representation similarity analysis

fMRI and EEG puzzle-induced brain activity were transformed to a common representational “puzzle space” using RSA (Kriegeskorte et al., 2008). RSA aims to model the similarity structure of brain activation patterns in response to differing tasks, conditions, or stimuli (**Figure 1**). Specifically, we estimated (dis)similarity between brain responses evoked by the 36 LST puzzles using the squared Euclidean distance. Representational dissimilarity matrices (RDMs), generated in both the fMRI and EEG datasets, were compared to a cognitive model, as well as each other (described below).

#### fMRI dataset

fMRI-RDMs were generated for 376 brain regions using the Glasser cortical parcellation (Glasser et al., 2016), and the Melbourne 3T S1 subcortex parcellation (Tian et al., 2020). For each participant and brain region, fMRI GLM coefficients (*z*-scored) for each of the 36 LST puzzles were organised into a puzzle-by-vertex (or voxel, in subcortex) activity pattern matrix. The RDM was then calculated as the puzzle-by-puzzle (36 x 36) squared Euclidean distance. Additionally, the resulting RDMs were averaged into thirteen previously established functional brain networks (Ji et al., 2019): auditory, cingulo-opercular, default, dorsal-Attention, frontoparietal, language, orbito-Affective, posterior-multimodal, somatomotor, subcortical, ventral-multimodal, and visual brain networks.

#### EEG dataset

For each participant and time point (1225 time points sampled at 250 Hz), preprocessed sensor-level EEG responses were organised into a puzzle-by-sensor activity pattern matrix. The RDM was then calculated as the puzzle-by-puzzle (36 x 36) squared Euclidean distance. This was performed for broadband signals and specific frequency ranges (theta, alpha, beta, and gamma, described previously). Following this, the RDMs were down-sampled into 50 sequential time segments of 100 ms each.

#### Relational complexity model

The RDM coding for the cognitive construct of relational complexity (Halford et al., 1998) is shown in supplementary **Figure 1,** a simplified depiction is illustrated in **Figure 1B**. According to this model, puzzles within the same complexity condition (Binary, Ternary, or Quaternary) evoke *similar* brain activation patterns, regardless of the specific visual configuration of stimuli within the puzzle (e.g., squares, triangles, circles and crosses). This model also predicts that puzzles from different complexity conditions will elicit *dissimilar* activation patterns in a graded fashion, with Binary (easy puzzles) and Quaternary (difficult puzzles) conditions being maximally distant. Therefore, brain regions or time periods exhibiting high correspondence to this cognitive model would provide evidence that the underlying neural representations are tracking relational complexity demand rather than other factors (e.g., stimulus arrangement).

#### Cognitive effort model

In a similar approach to Chiou et al. (2024), we defined a data-driven ‘cognitive effort’ model. The model was generated using behavioural performance from both datasets. Specifically, the Euclidian distance was calculated between each puzzle’s error rate (1 – accuracy) and response time (averaged across participants). The complete model used in the RSA is shown in supplementary **Figure 1** and illustrated in **Figure 1B**. In this model, problems that participants found easier or harder are clustered together regardless of complexity level.

#### fMRI-EEG fusion analysis

We used commonality analysis (Hebart et al., 2018) to determine how much of the shared variance between EEG and fMRI RDMs could be explained by our RC and cognitive effort models. The commonality coefficient compares two semi-partial correlation coefficient (Spearman correlation). The first specifies the proportion of variance shared between EEG and fMRI when partialling out all model variables except the model of interest (e.g., the RC model) from fMRI. The second reflects the proportion of shared variance between EEG and fMRI after partialling out all model variables (including the variable of interest). The commonality coefficient is the difference between both coefficients of determination (*R^2^*). We performed this analysis for each brain network and time point, tracking the unique contributions of each model across space and time. Formally, for time *t* and network *j* the commonality coefficient is defined as:

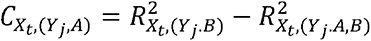

where *X* indicates EEG, *Y* indicates fMRI, *A* reflects the RC model, and *B* reflects the cognitive effort model.

### Follow-up fMRI frontoparietal and EEG whole-brain sensor activity

As a supplementary analysis, we examined how activity in the FPN varied across the three complexity conditions. To do so, we averaged the GLM beta coefficients across each condition for each participant within each FPN region. These values were averaged across regions to generate a single value per participant, per condition.

For the spectral characteristics of the EEG data, we computed the power spectral density (PSD) using Welch’s method within the data-driven beta RSA time window of interest, spanning 2.1 to 4.2 seconds post-stimulus onset. The PSD was calculated using segments of 500 samples with a 50% overlap. We applied a frequency range of interest from 2 to 45 Hz. Each participant’s PSD were averaged over trials and channels to obtain a whole-brain sensor level measure of power for each complexity condition.

The periodic and aperiodic spectral parameterisation algorithm (version 2.0.0rc2) was used to parameterise neural power spectra (Donoghue et al., 2020). Settings for the algorithm were set as: peak width limits: (0.5, 12.0); max number of peaks: inf; minimum peak height: 0.0; peak threshold: 2.0; and aperiodic mode: ’fixed’. Power spectra were parameterized across the frequency range 2.0 to 45.0 Hz. High model fits of the parametrised spectrum were consistent across complexity conditions: 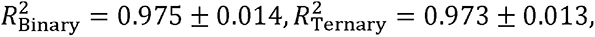 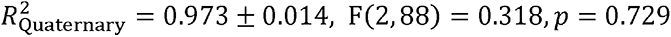. Using the previously defined fixed beta (13-30 Hz) frequency range we extracted the periodic peak frequency for each subject across conditions from the modelled spectrum. Five subjects were excluded as a peak could not be defined across all conditions.

### Statistical analysis

Task performance (accuracy and response time) in each dataset was assessed using a one-way repeated measures ANOVA. To compare performance between the datasets, Spearman correlations were conducted on the participant-averaged puzzle level performance (accuracy and response time) between the two datasets.

The spatial (fMRI) and temporal (EEG) RDMs were compared with the RC and CE models RDM (**Figure 1B**) using Spearman rank correlation without tie correction (Nili et al., 2014). For each participant and each brain region, network, or epoch, the vectorized upper triangle of each empirical RDM was correlated with the vectorized upper triangle of the model RDM. These correlation values - or coding strengths - underwent normalisation via the Fisher z- transformation, and were entered into a one-sample *t*-test (Vallat, 2018). All three analyses were multiple comparisons corrected using the False Discovery Rate (FDR), a = 0.05 (fMRI n_regions_ = 376, n_networks_ = 13, EEG n_epochs_ = 50) (Benjamini and Yekutieli, 2001).

Statistical significance in the fMRI-EEG fusion analysis was assessed using a permutation test within each time bin (N=5000, (Hebart et al., 2018)). For each permutation, we randomly shuffled the rows and columns of the EEG RDMs (Kriegeskorte et al., 2008) and recomputed the commonality coefficients, generating a null distribution of coefficient time courses for each network. P-values were calculated as the proportion of permutations that exceeded the empirical commonality coefficient. These p-values were subsequently corrected for multiple comparisons using FDR.

Finally, an exploratory analysis was conducted to relate individual differences in brain representations and LST performance. Specifically, a Spearman correlation was performed between the RC model coding strength and LST accuracy. For the EEG data, each individual’s peak coding value was used. Given the exploratory nature of this analysis it was only performed in networks and frequency bands that demonstrated high coding strengths in the initial analyses (four networks and two frequency bands). A one-way repeated measures ANOVA was used to assess the follow-up brain activation measures of the frontoparietal network and the beta band power.

### Code and data availability

The fMRI data analysed in this study are openly available (https://espace.library.uq.edu.au/view/UQ:734743). The EEG data will be made available to researchers following the execution of a data transfer agreement with QIMR Berghofer. Code to perform the analyses is available on Github (released upon publication).

## Results

### Effect of relational complexity on accuracy and response time

Participants completed relational reasoning problems at three complexity levels (**Figure 2**). As shown in **Figure 2A** (top), results from both fMRI and EEG datasets demonstrated the characteristic degradation in performance accuracy as complexity increased: Binary (Mean_fMRI_ = 96.66%, Mean_EEG_ = 95.00%), Ternary (Mean_fMRI_ = 87.02%, Mean_EEG_ = 80.69%), and Quaternary (Mean_fMRI_ = 66.25%, Mean_EEG_ = 56.30%); F(2, 78)_fMRI_ = 80.61, p < 0.0001, and F(2, 88)_EEG_ = 101.60, p < 0.0001. Correspondingly, response times **Figure 2A** (bottom), increased with complexity: Binary (Mean_fMRI_ = 0.74 ms, Mean_EEG_ = 0.61 ms), Ternary (Mean_fMRI_ = 0.81 ms, Mean_EEG_ = 0.78 ms), and Quaternary (Mean_fMRI_ = 0.93 ms, Mean_EEG_ = 0.87 ms); F_fMRI_(2, 78) = 51.09, p < 0.0001 and F(2, 88)_EEG_ = 108.00, p < 0.0001. To assess consistency across datasets, puzzle performance was averaged across participants in each dataset and correlated. Both accuracy (rs = 0.95, p < 0.001) and response time (rs = 0.83, p < 0.001) demonstrated a high correspondence between datasets (**Figure 2B**).

**Figure 2.**
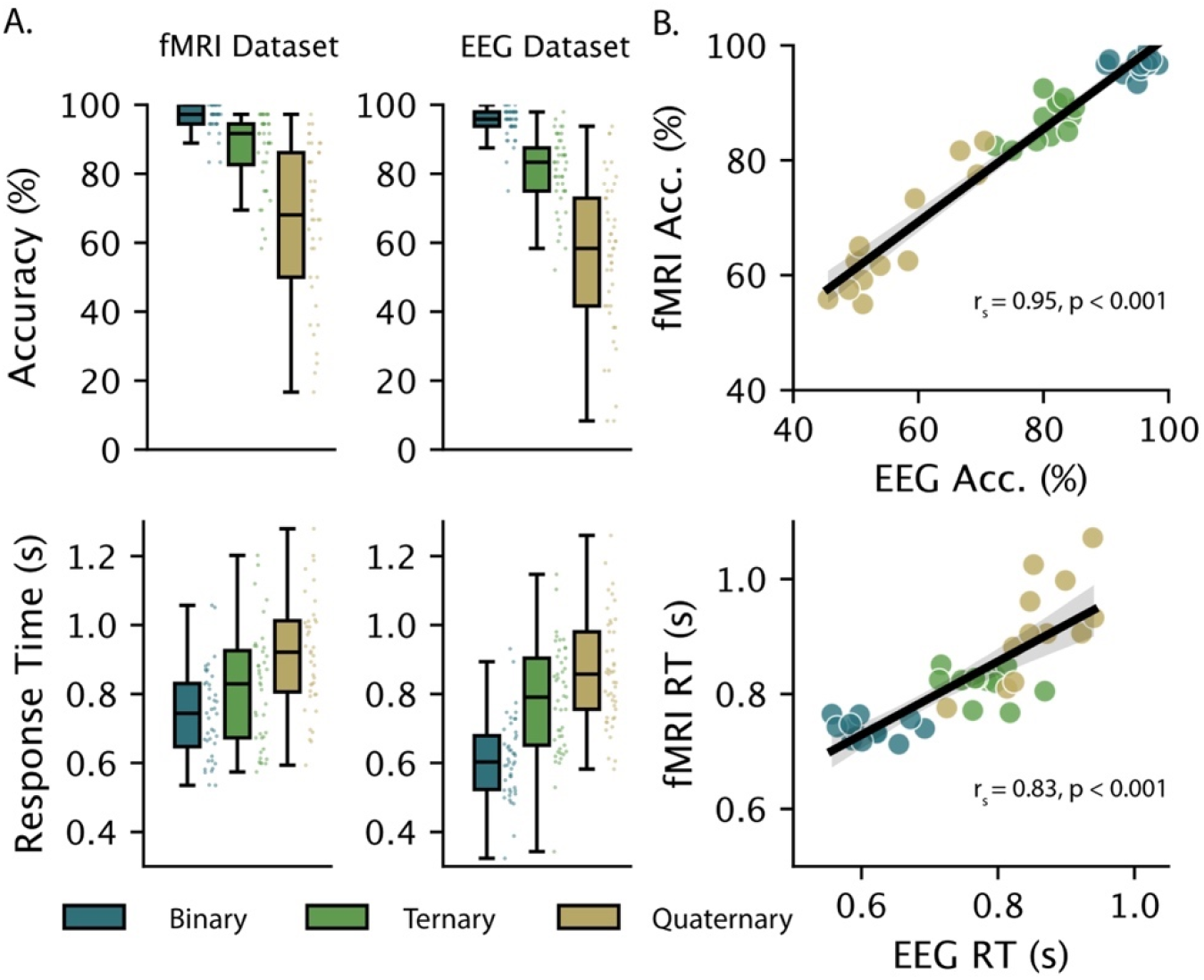
*Behavioural performance in the Latin Square Task*. **A.** Results replicated the expected degradation in accuracy (top panel) and delay in response time (bottom panel) associated with increased relational complexity across both fMRI (n=40) and EEG (n=45) datasets. **B.** Participant performance on the original 36 puzzles showed a positive correlation between fMRI and EEG datasets for both accuracy (top panel, rs = 0.95, p < 0.001) and reaction time (bottom panel, rs = 0.83, p < 0.001).

### Spatial and temporal emergence of relational complexity and cognitive effort

We conducted an RSA comparing models discerning relational complexity and cognitive effort (**Figure 1B**) to representations derived from puzzle-evoked fMRI and EEG data. In the fMRI data, analyses were conducted at the region (**Figure 3A and 3C**, *left*) and network level (**Figure 3A and 3C**, *right*). Surprisingly, we observed strong relational complexity coding expressed across the whole brain. Of the 376 possible brain regions tested, 367 demonstrated significant correspondence to the model (*p_FDR_*< 0.05). However, as expected, peak representational values were observed in prefrontal and parietal cortices: the superior premotor subdivision (6a), superior parietal cortex (7PL, 7Pm, 7Am, VIP, AIP, MIP), inferior parietal cortex (IP2, PGp), dorsal visual transitional area (DVT), anterior ventral insular area (AVI), medial prefrontal cortex (8bm) and dorsolateral prefrontal cortex (8C, i6-8). Accordingly, transmodal brain networks associated with goal-directed behaviour and higher-order cognition (Cole and Schneider, 2007; Duncan, 2010; Assem et al., 2020) demonstrated the strongest similarity to the RC model. Similar to the findings of the region-wise analysis, all networks showed statistical similarity to the RC model (all networks survived *p_FDR_* < 0.05). The CE model also exhibited broad representation across brain regions and networks (365 regions and all networks survived *pFDR* < 0.05), though with reduced similarity compared to the RC model.

**Figure 3.**
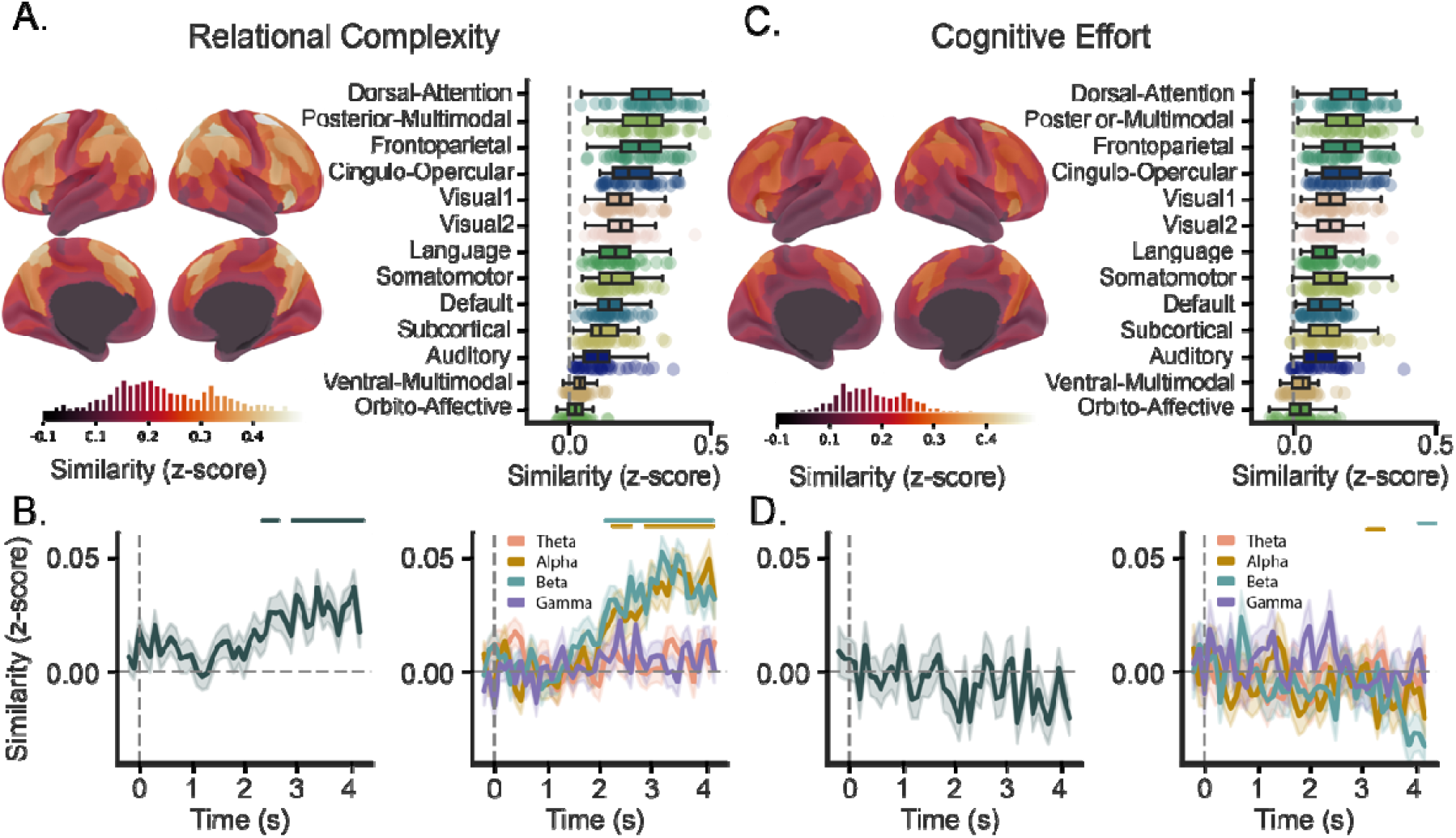
*Spatial and temporal signatures of relational complexity (RC) and cognitive effort (CE) derived from fMRI and EEG data*. Group averaged brain-wide correlations between fMRI-informed representations and the RC (**A**) and CE (**C**) models, colour bar indicates a transition from low (dark) to high (bright) correspondence. Underneath a histogram displays the frequency ROI correlations mapped onto brain surface renderings (Glasser et al., 2016; Tian et al., 2020). *Right*, fMRI results when down sampled into canonical brain networks (Ji et al., 2019), ordered by effect size. Group averaged correlation between EEG-informed representations and the RC (**B**) and CE (**D**) models across time. *Right,* frequency band derived EEG representations and the cognitive models across time, theta (4-8 Hz, pink), alpha (8-12 Hz, gold), beta (13-30 Hz, teal), and gamma (30-45 Hz, purple). Coloured line(s) above indicate statistically significant time bins (*p_FDR_* < 0.05). Confidence intervals indicate standard error of the mean. All correlation values are Fisher-Z transformed.

To identify when the relational complexity construct emerged in time, the model was correlated across the RDMs derived from the EEG broadband timeseries. The similarity timeseries (**Figure 3B**, *left*) revealed that RC representations emerged during the 2.5 – 4.1 seconds puzzle-stimuli window (*p_FDR_* < 0.05). At the group level, the temporal trend of the representation built over time, peaking at 4.1 s (peak correlation z = 0.037). In contrast to RC, the CE model showed no significant representation in the EEG broadband timeseries (**Figure 3D**, *left*).

As a follow up analysis, we repeated the analysis in four canonical frequency bands (**Figure 3B and 3D**, *right*). Interestingly, we only found significant RC model coding in alpha and beta frequencies (alpha 2.3 – 4.2 s, beta 2.1 – 4.2 s, *p_FDR_* < 0.05). Both frequencies mirrored the temporal trend of the broadband signal, peaking at 4.1 s (peak correlation, z = 0.049) and 3.2 s (peak correlation z = 0.052) for alpha and beta, respectively. While the CE model was also represented in alpha and beta frequencies, these representations were transient and occurred at discrete windows (alpha 3.3 s, beta 4.2 s, *p_FDR_* < 0.05). Together, our results suggest that RC is i) most strongly represented in brain networks supporting goal-directed external behaviour (e.g., dorsal attention and frontoparietal networks), ii) emerges late in the trial period, and iii) is specifically related to alpha and beta frequency bands. Notably, although CE showed widespread but weaker spatial representation, its brief frequency-specific temporal representation suggest that during the LST the brain primarily organises neural resources according to RC rather than general task difficulty.

### Cross-modal fusion analysis of time and frequency-resolved patterns of network representations

Next, we asked when the representational structure in a given brain network (determined from fMRI) corresponds with the representations from the time-resolved EEG signal. Using the shared representation between modalities, we further wanted to determine how much RC accounted for the explained variance in the representation. To do so, we performed a model-based fMRI-EEG fusion commonality analysis (**Figure 4a**) (Cichy et al., 2014; Hebart et al., 2018). This approach also allowed us to compare the contribution of the complementary CE model derived using participant performance. In contrast to RC, the CE model proposes that the information content coded in the representation can be retrospectively explained by the challenge in mapping the underlying abstraction due to a confluence of puzzle-specific properties (such as the number or configuration of items). Conversely, RC ignores puzzle-specific properties and proposes that the information in the signal codes for the dimensionality of the abstraction. Although both models aimed to capture different theoretical constructs (effort vs. complexity), they showed a high correlation (rs = 0.81, p < 0.001). Based on the prior results (**Figure 3B**), we narrowed our focus to the alpha and beta frequency bands.

**Figure 4.**
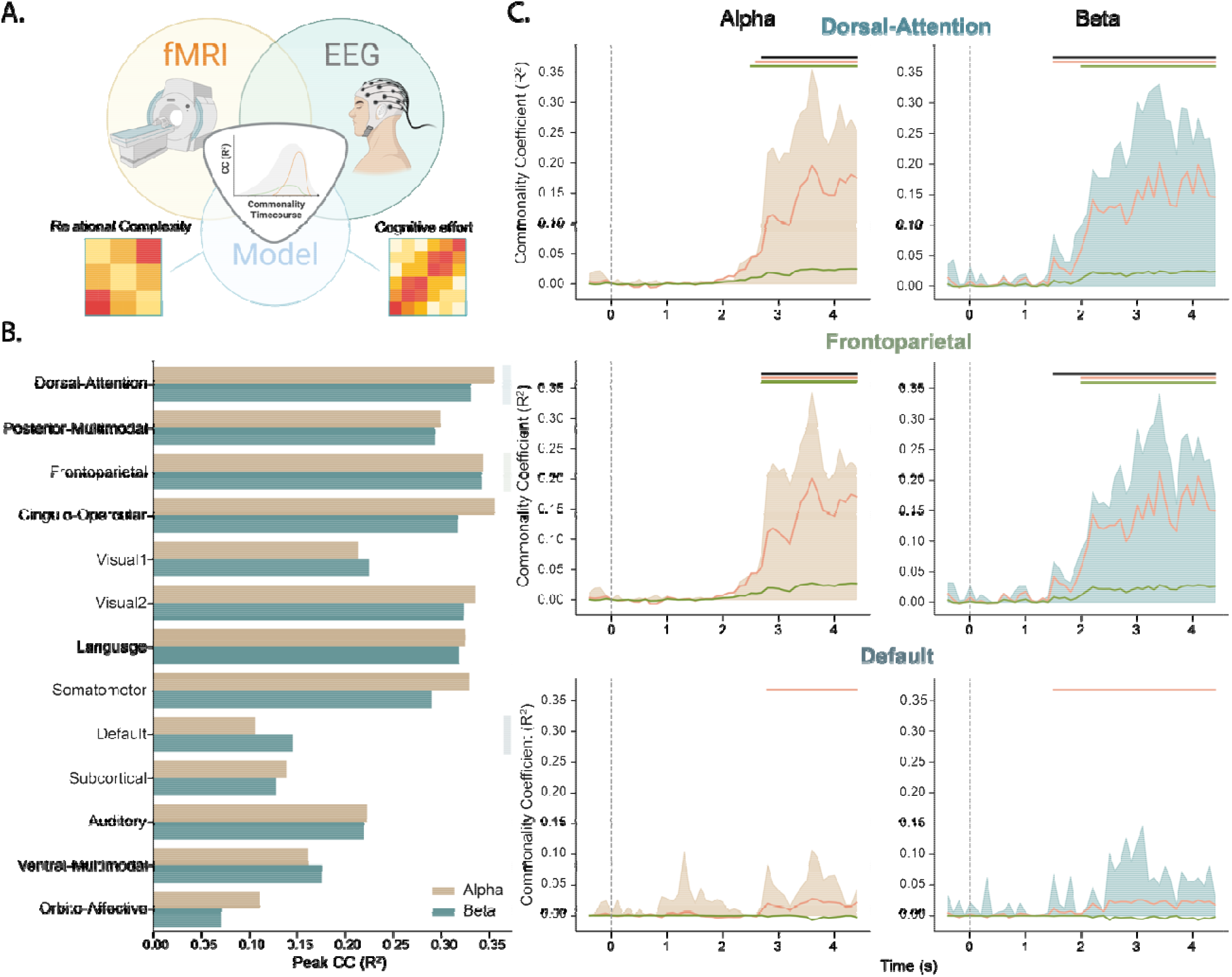
*Model based EEG-fMRI fusion-analysis*. **A.** Commonality analysis schematic. The commonality coefficient (CC) reflects the variance shared between the fMRI and EEG signal that can be attributed to specific task models (RC and CE). **B.** Peak correlations between averaged fMRI and EEG data for each network within the alpha (gold) and beta frequencies (teal). **C.** Commonality coefficient timecourses for the RC model (orange line) and CE model (green line) in three exemplar brain networks (dorsal-attention; top, frontoparietal; middle, default mode; bottom) in both the alpha (right) and beta (left) frequency bands. The shaded area indicates the proportion of variance shared between EEG and fMRI data over time.

The peak commonality coefficients from the fMRI-EEG fusion results in alpha and beta frequency bands are shown in **Figure 4B**. Most networks exhibit a similar peak onset time at 3.8 s in the alpha, while beta appeared to emerge 200 ms earlier at 3.6 s. Higher-order dorsal attention (peak alpha R^2^ = 0.35, beta R^2^ = 0.33), cingulo-opercular (peak alpha R^2^ = 0.36, beta R^2^ = 0.32) and frontoparietal (peak alpha R^2^ = 0.34, beta R^2^ = 0.34) networks shared the highest peak similarity between imaging modalities. The top networks showing the highest peak representation were consistent across both frequency-resolved timeseries. To further analyse the variance explained by our proposed cognitive models (**Figure 4B**), we selected the dorsal- attention (top) and frontoparietal (middle) networks, shown in **Figure 4C**, since they shared the highest cross-modal commonality. We also selected the default mode network (bottom), a transmodal network known to be anti-correlated in response to RC demands (Hearne et al., 2015) that showed relatively low peak fusion similarity (**Figure 4B**). The variance captured between modalities, **Figure 4C**, alpha (left) and beta (right) is shown by the shaded area. A shared representation in the beta band for both dorsal attention (alpha 2.3 – 4.2 s, beta 2.1 – 4.2 s, *p_FDR_* < 0.05) and frontoparietal networks (alpha 2.3 – 4.2 s, beta 2.1 – 4.2 s, *p_FDR_*< 0.05) preceded and maintained longer representation than the alpha band. The default mode network did not show any period with a significant commonality coefficient.

The commonality coefficients of the RC and CE models showed a similar trend to the overall correlation between the two imaging modalities over time. After 2.5 seconds post stimuli onsets both models exhibited higher commonality coefficients than would be expected by chance. Of note, between both models, RC is shown to account for most of the variance of the shared representational structure. The cognitive effort model was not present in the default mode network. However, the RC model was detected in the signal despite the low shared variance between EEG and fMRI.

### Peak representation of RC and individual task performance

In a final exploratory analysis, we correlated individual participant task accuracy with RC i) model coding strength across brain networks and ii) peak oscillatory model coding strength in the alpha and beta frequency. We observed a negative correlation between task accuracy and RC model coding strength in the frontoparietal (rs = -0.32, p = 0.045) and cingulo-opercular networks (rs = -0.33, p = 0.038, **Figure 5A**). Focusing on the Frontoparietal network (**Figure 5C**, *left*), we observed an increase in network activation as RC increased: Binary (Mean = - 0.027), Ternary (Mean = 0.547) and Quaternary (Mean = 0.786); F(2, 78) = 117.00, p < 0.001.

**Figure 5.**
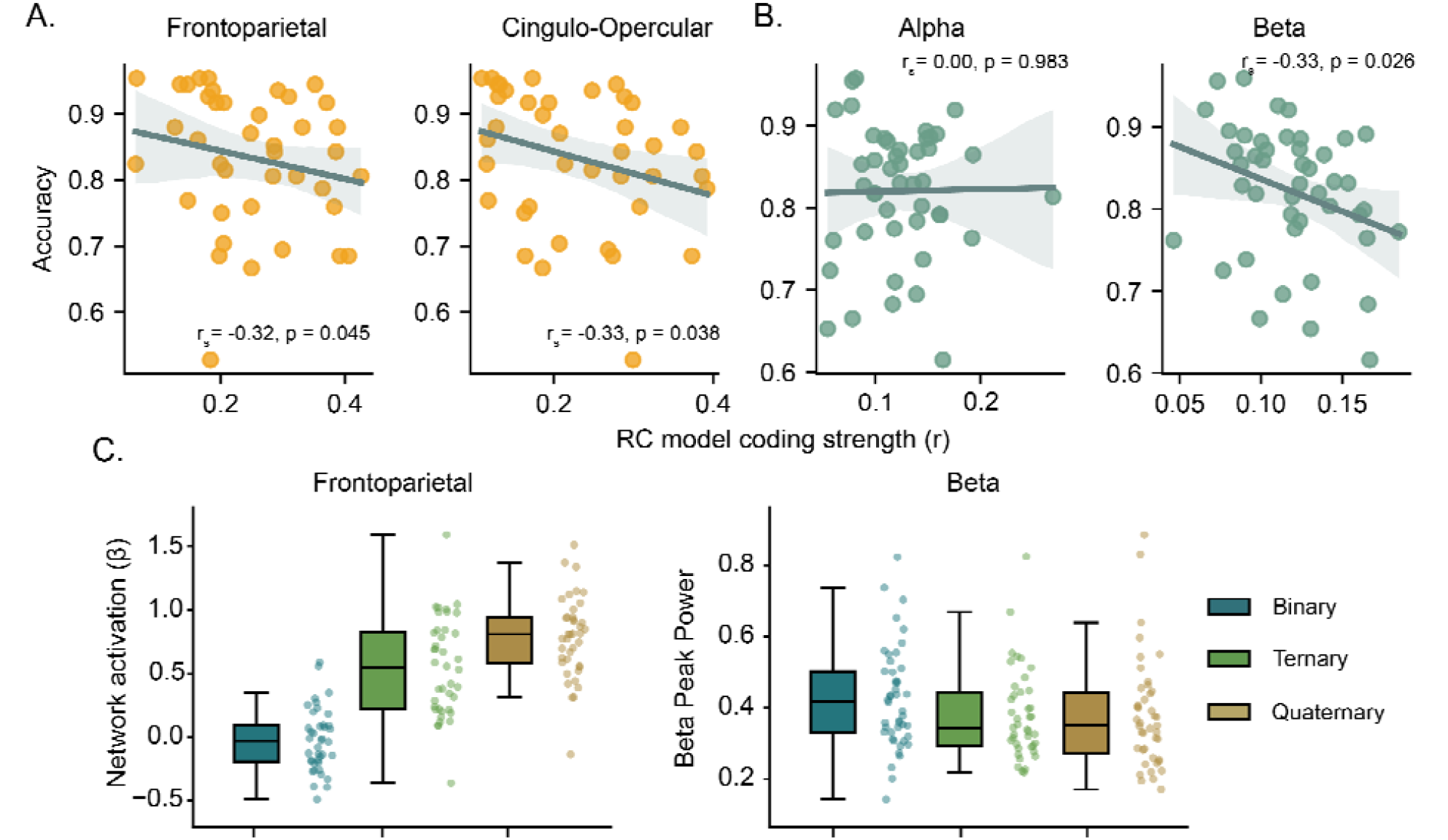
Correlation between participant accuracy (y-axis) and RC model coding strength (x- axis) with corresponding brain activation across complexity levels. **A.** fMRI RC model coding strengths for the frontoparietal (left) and cingulo-opercular (right) networks. **B.** EEG frequency derived peak RC model coding strengths in the alpha (8 – 12 Hz, left) and beta (13 – 30 Hz, right) bands. rs represents Spearman’s rank correlation coefficient; p-values are uncorrected. **C.** Left. Frontoparietal activation (β coefficients) across complexity levels. *Right.* Periodic peak beta power calculated across all EEG electrodes in each complexity condition during the 2.1s to 4.2 s time window.

In the EEG dataset, task accuracy also had a negative correlation with the temporal peak representation in the beta oscillatory band (rs = -0.33, p = 0.026), while no relationship was found in the alpha band (rs = 0.00, p = 0.983, **Figure 5B**). Contrary to RC, we did not find evidence to suggest that the spatial or temporal CE neural representations were linked to performance (**Supplementary** Figure 2). Further exploration of sensor-level activity during the RC coding window of the beta band (**Figure 3B**, 2.1 s to 4.2 s) showed a decrease in peak beta power with RC: Binary (Mean = 0.437), Ternary (Mean = 0.380) and Quaternary (Mean = 0.388); F(2, 80) = 9.68, p < 0.001 (**Figure 5C**, *right*). While the reported correlations do not survive appropriate correction for multiple comparisons, the reliable pattern of results across key cognitive control networks and modalities supports the genuine association between task-accuracy and the strength of RC brain representations.

## Discussion

In this multimodal study, we assessed the brain basis of RC representations in the context of a visuospatial reasoning task. Using representational similarity analysis, we showed that RC i supported by a widespread network of regions, with core contributions from functional brain networks associated with external goal-directed control. Results from the analysis of macroscale temporal dynamics showed that trial-based RC representations are supported by alpha and beta brain rhythms and emerged gradually, with peak correspondence in the 2.5 – 4.1 second post-stimuli onset window. The RC model accounted for more variability in the correlation between EEG-fMRI task representations than the CE model. Moreover, individual RC coding strength in specific brain networks and frequencies correlated with task performance. This set of results advances our understanding of the spatiotemporal dynamics underpinning complex reasoning, importantly unveiling that RC is coded in the representation of problems. Considering the strong association between RC and intelligence (Hansell et al., 2015), our findings showcase how distributed neural processes progressively integrate relations to orient intelligent behaviour.

Our fMRI findings build on prior knowledge to show that RC is distributed throughout the brain. Previous lesion and neuroimaging work have demonstrated a key role of frontal (Christoff et al., 2001; Bunge et al., 2005; Wendelken et al., 2008) and frontoparietal circuits (Vendetti and Bunge, 2014; Wendelken et al., 2016) in supporting relational processes. However, several studies have expanded this view by showing that changes in task complexity are supported by local modulations in brain activity and connectivity in brain network regions beyond the frontal and parietal cortices (Hearne et al., 2017). For example, Cocchi et al. (2014) demonstrated that progressively increasing problem complexity also resulted in a proportional modulation of brain activity and task-based connectivity between core nodes of the cingulo-opercular network. These observations are in line with network-centric views of cognition and relational integration (Cole and Schneider, 2007; Morin et al., 2022), positioning frontoparietal hubs and brain-wide connectivity as central to orchestrating goal-directed cognitive control (Cole et al., 2013). Our findings extend this work by demonstrating, for the first time, that relational complexity is mapped in the neural representation of frontoparietal, dorsal-attention, and cingulo-opercular networks. Visual networks also had high model correspondence, likely reflecting the interaction between the sensory cortex involved in the task and the transmodal ‘multiple demand’ cortex (Cocchi et al., 2014; Hearne et al., 2017). On the other hand, the default mode network in the fusion analysis did not show a strong association with the RC model, suggesting that previously observed complexity-driven changes in DMN activity and connectivity (Hearne et al., 2015) may be more related to the engagement of task-positive networks rather than the coding of RC.

In a related fMRI-RSA study examining basic arithmetic problems, RC was held at a consistent binary level (addition problems), while altering factors such as digit magnitude, carryover operations, and context (easy vs. hard) showed neural MDN patterns scaled with cognitive effort (Wen and Egner, 2023). Consistent with previous findings we were also able to demonstrate that part of the neural information stored in higher-order regions reflected the behavioural effort taken to solve the problems (Chiou et al., 2024). Despite the high correlation between RC and CE models, the neural information content in the higher-order networks in our study predominantly coded for the dimensionality of the abstraction as proposed to RC theory, rather than trial-specific puzzle factors associated with cognitive effort.

Coherent oscillatory activity is a fundamental element in computational theories of relational integration (Knowlton et al., 2012; Fries, 2015). Our results suggest that the widespread neural representations of RC are supported by alpha and beta oscillatory brain rhythms, which emerge after the first two seconds of stimulus onset. These frequency-resolved alpha and beta results complement our fMRI findings by showing the temporal dynamics supporting the establishment of RC representation in task-relevant dorsal attention, frontoparietal, and cingulo-opercular networks. RC representation emerged following the initial computation of the perceived properties of a stimulus (shape, colour; (Cichy et al., 2014)) and mirrors the timeframe observed to implement task-relevant rules (Hebart et al., 2018). The involvement of neural activity in the alpha frequency range is consistent with findings suggesting that frontal and parietal alpha activity support logical reasoning (Brzezicka et al., 2011; Brzezicka et al., 2017; Heldmann et al., 2024). Accordingly, frontoparietal alpha and beta-rhythms have been associated with performance on matrix reasoning tasks (Ociepka et al., 2023; Penhale et al., 2024). In contrast, we found no evidence for theta or low gamma (30–45 Hz) representations that correspond to either RC or CE models. Although high gamma activity (>80 Hz) has been strongly associated with higher-order cognitive processing (Honkanen et al., 2015; Lundqvist et al., 2018), it is unclear whether these patterns are absent due to methodological limitations (e.g., lack of high-frequency neural signals with scalp-based EEG).

Based on prior literature, we can speculate that alpha-band activity may serve to suppress irrelevant sensory inputs (Foxe and Snyder, 2011; Gray et al., 2015) or competing relational mappings (Benedek et al., 2014), thereby facilitating the focus on task-relevant information. In contrast, beta-band oscillations are likely to support the sustained activation of task-specific neural activity (Buschman et al., 2012), facilitating the maintenance of relational representations as they are progressively integrated. Other cognitive experiments in particular working memory, attention and rule-switching tasks indicate these high-order cognitive operations decrease alpha and beta power in response to increased demands (Scharinger et al., 2017; Williams et al., 2019; Kaiser et al., 2023). However, this attenuated activity appears to facilitate an increase in band-specific synchronisation between brain regions (Sauseng et al., 2005; Palva et al., 2010; Buschman et al., 2012). While our study cannot disentangle the exact contribution of network-specific alpha and beta rhythms to support RC representations due to the constraint of the LST design, the present work provides a solid rationale to further investigate these neural correlates.

In a final exploratory analysis, we found that individual differences in task performance were inversely related to RC model coding within the beta oscillatory band (EEG dataset) and the frontoparietal and cingulo-opercular networks (fMRI dataset). In other words, participants with lower accuracy (‘poor performers’) produced brain representation patterns that corresponded more to the RC model. The mechanism behind this unintuitive result is unclear, but it would suggest low performers tend to have more clearly delineated boundaries between the brain representations of different complexity levels (and vice versa regarding better performers). One interpretation of this finding is that high performers require less attention-related resources on more difficult problems, due to more efficient processing (Neubauer and Fink, 2009). However, our CE model - presumably a better test of this hypothesis – showed no such relationship with participant task performance. Cognitive effort, relational complexity and attentional demands are difficult to tease apart experimentally but additional behavioural measurements or physiological measures (e.g., pupillometry) would likely be beneficial in future studies.

Given the existence of theoretical accounts of relational reasoning (Knowlton et al., 2012), as well as neural network LST solvers (Ito et al., 2024), a key future research direction is to leverage the RSA framework to test specific model-based hypotheses regarding RC. In the version of the LST we implemented in this study, the puzzles were designed to minimise the loading of working memory demands across complexity levels, reducing multiple processing steps to isolate RC. However, other variations of RC tasks may allow for alternative processing strategies, such as segmentation or chunking, that can overcome trial-specific complexity based on problem attributes (Hartung et al., 2022). This opens up opportunities to explore more nuanced questions of relational integration by examining the trade-offs involved in building complex representations across a diverse range of visuospatial, semantic, and analogical tasks (Halford et al., 1998), and addressing the broader question of whether RC is represented across a domain-general set of problems. A model comparison approach (Schutt et al., 2023), as demonstrated in recent studies (Daws et al., 2020; Chiou et al., 2024), would help to directly contrast different models of reasoning. One insight we gained from this method – when combined with a fusion analysis (Hebart et al., 2018) – was that RC captured the majority of variance shared between EEG and fMRI when contrasted with a cognitive effort model. It is worth noting however, that different participant cohorts were used in the multimodal fusion analysis, potentially weakening the correspondence between the fMRI and EEG datasets.

In conclusion, our study provides a novel characterisation of the spatial and temporal brain processes underpinning representations of complexity during a relational reasoning task. Our results highlight the role of domain-general functional networks and suggest a key role of alpha and beta oscillatory patterns in the service of human relational reasoning. The current results advance our understanding of the neural mechanisms of reasoning and facilitate the establishment of formal generative models of RC.

## Acknowledgments

C.R is supported by an Australian Government Research Training Program Scholarship and a Living Stipend through the penurious administration of the University of Queensland. C.R is also supported by a QIMR administered Living Stipend extension funded by L.J.H. **Figure 1** and Figure 4A was created with BioRender.com. This work was supported by the Australian NHMRC (LC: GN2001283 and GNT2027597, LJH: APP1194070).

## Author contributions

L.J.H. and C.R conceptualised the research goals; L.J.H. supervised the investigation, collected and pre-processed dataset 1 and assisted with the formal data analysis. C.R collected and pre- processed dataset 2 formally analysed the data and wrote the first draft. All authors reviewed and edited the manuscript.

