## Supplemental Figures 1 and 2 for "Relational integration demands are tracked by temporally delayed neural representations in alpha and beta rhythms within higher-order cortical networks"

**Supplementary**

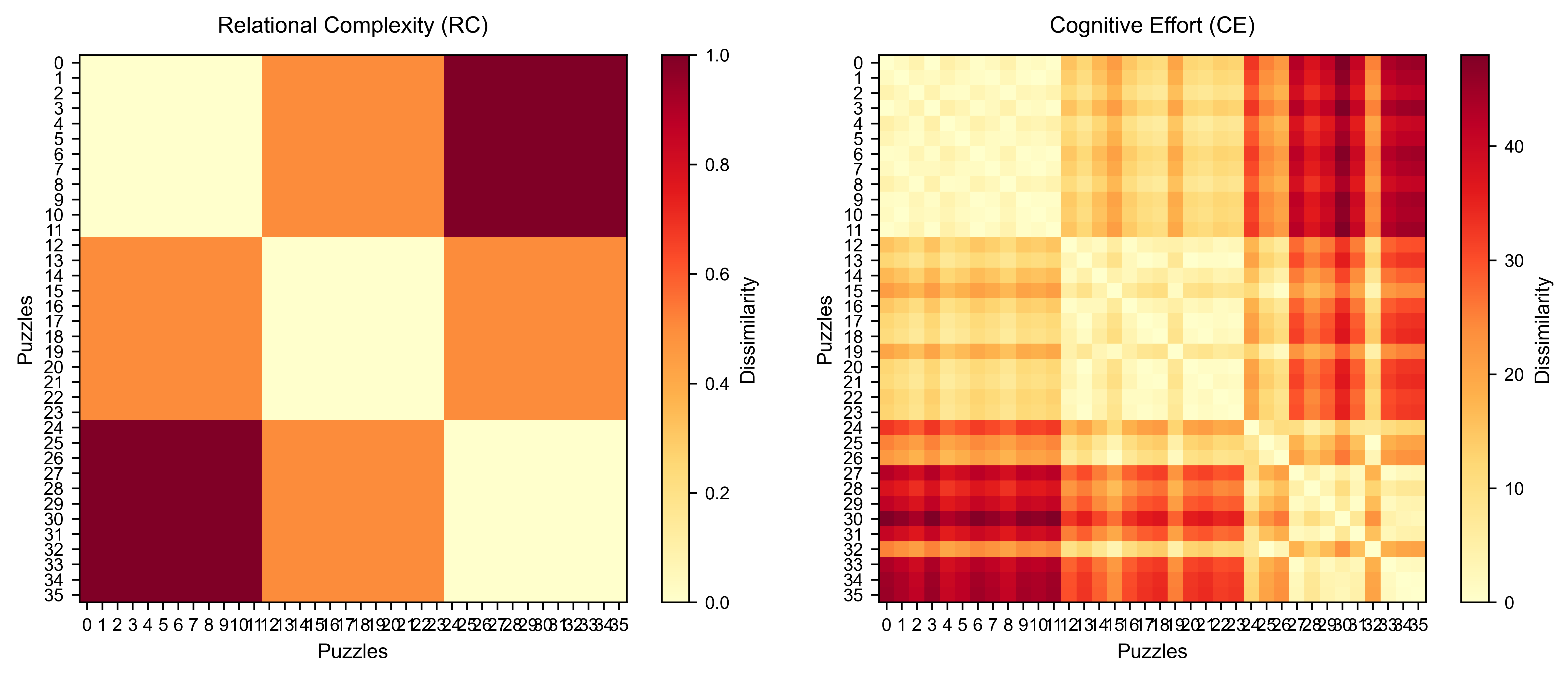

**Supplementary Figure S1**: *Representational dissimilarity matrices (RDMs) for relational complexity (RC) and cognitive effort (CE) models.* The RC model (*left*) defines puzzle dissimilarity based on the number of relations that must be integrated to reach a solution, with binary, ternary, and quaternary puzzles assigned values of 0.0, 0.5, and 1.0, respectively. The CE model (*right*) is a data-driven model, derived by calculating the Euclidean distance between puzzles in a 2D feature space of error rate (1 – accuracy) and response time. Both matrices show pairwise dissimilarity between puzzles, where each row and column represents an individual puzzle. In both cases, cooler colours (yellow) indicate greater similarity, while warmer colours (dark red) indicate greater dissimilarity. Simplified cartoon illustrations are depicted in **Figures 1B** and **4A**.

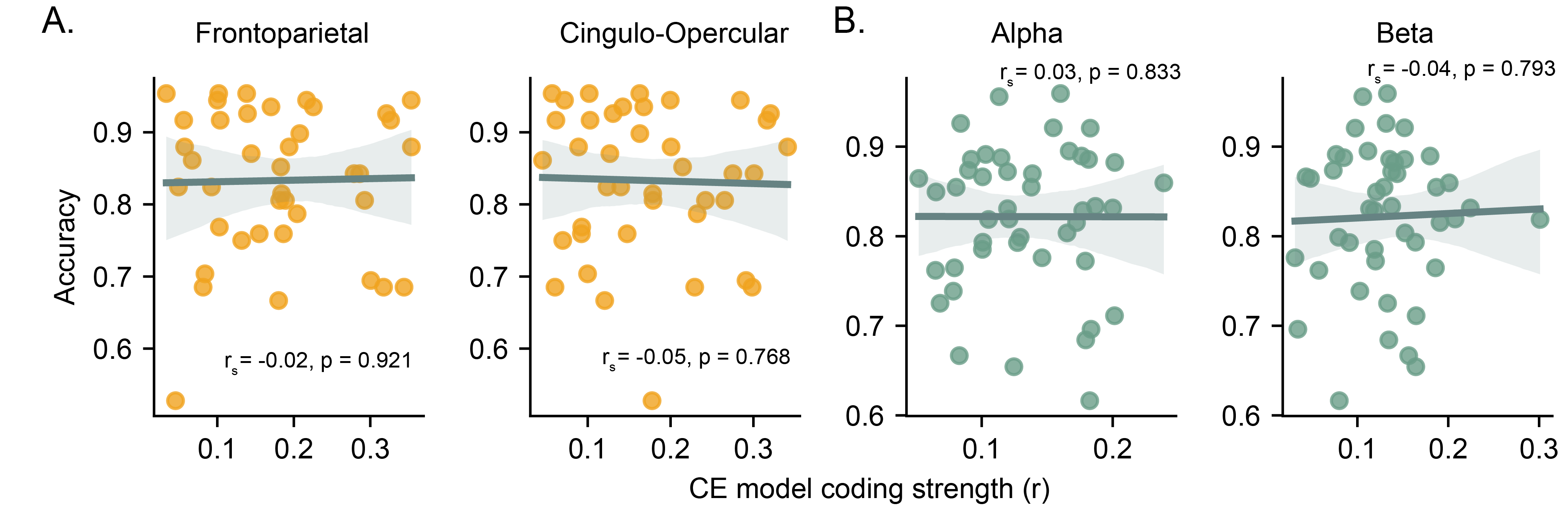

**Supplementary Figure S2**: *Correlation between participant accuracy (y-axis) and CE model coding strength (x-axis)*. **A.** fMRI CE model coding strengths for the frontoparietal (left) and cingulo-opercular (right) networks. **B.** EEG frequency derived peak CE model coding strengths in the alpha (8 – 12 Hz, left) and beta (13 – 30 Hz, right) bands. rs represents Spearman's rank correlation coefficient; p-values are uncorrected.
